# Metadynamics Simulations meet Ligand Design for the Reversible Inhibition of Human Peroxiredoxin 5

**DOI:** 10.1101/2025.04.25.650576

**Authors:** Laura Troussicot, Florian E.C. Blanc, Yoann Pascal, Sebastien Vidal, Jean-Marc Lancelin, Florence Guillière

## Abstract

Using insights from Funnel Metadynamics, a molecular dynamics protocol that provides a detailed representation of protein-ligand interactions, we investigated how a single heavy-atom modification can enhance the activity of an initial hit against human peroxiredoxin 5. This improvement was validated by NMR experiments and enzyme inhibition assays. Our results illustrate how molecular dynamics simulations offer a rational framework for designing ligands with improved properties starting from low-affinity but selective hits with minimal structural modifications.

**TOC Graphic:** 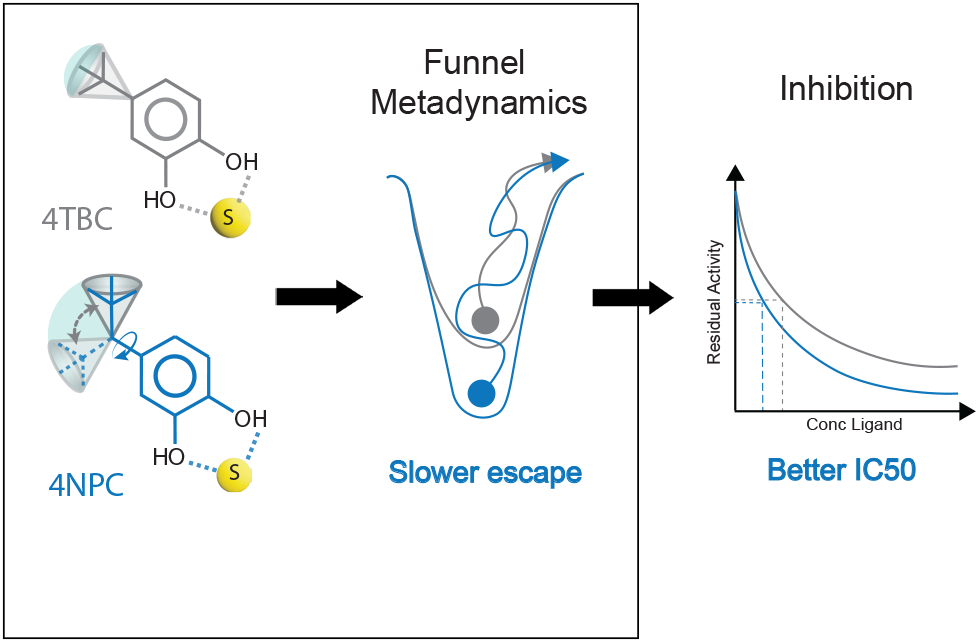

## Introduction

The design of ligands to modulate the activity of biological receptors is central to drug discovery.^1,2^ In fragment-based approaches, low-affinity but selective simple molecules are optimized through chemical modifications until they bind to the target with maximum lasting efficiency.^3,4^ In this study, we highlight how quantitative-accuracy Molecular Dynamics simulations can enable the rational design of an improved inhibitor against an enzyme relevant for Human health.

The thermodynamics of ligand-receptor binding can be studied at the macroscopic level by *in vitro* biophysical methods (e.g., NMR, enzyme kinetics) or computationally by Molecular Dynamics (MD) simulations. A key interest of MD is that it provides atomically detailed insight into the drug-target interaction. However, quantitative prediction of binding affinity by simulations remains challenging and requires computationally costly free energy calculation techniques.^5–10^ Among them, alchemical methods evaluate the free energy difference along artificial transformations in which the protein/ligand interactions are progressively turned on (or off).^11^

These methods can accurately predict the binding affinity, but their non-physical character makes it challenging to infer mechanistic information. Alternatively, geometric methods evaluate the binding affinity by collecting the change in free energy along geometric degrees of freedom that describe the binding process, such as the distance between the ligand and the binding site.^12^ This can provide more realistic molecular details than in the alchemical case. However, reaching convergence requires specialized protocols to control the translational, rotational, and conformational entropy of the ligand. Funnel metadynamics (FM) combines metadynamics, a popular geometrical free energy method,^13,14^ with a funnel-shaped restraining potential to focus the sampling on reversible binding events.^15^ This promotes the convergence of the binding free energy estimate. Previously, we highlighted the potential of FM for drug design by demonstrating how it can quantitatively predict binding affinities of enzymatic inhibitors.^16^ Since then, FM has been widely used for elucidating ligand binding modes of several biological systems. ^17,18^ The connection between the microscopic and macroscopic descriptions of binding provided by the FM free energy landscape offers a powerful way to engineer the affinity of ligands against a known receptor, rationalizing the optimization process. Here, we apply FM to the design and validation of an inhibitor of Human peroxiredoxin 5.

Human peroxiredoxin 5 (hPrx5) is a thiol-active peroxidase that catalyzes the decomposition of hydroperoxides, and is part of the peroxiredoxin superfamily.^19,20^ Peroxiredoxins are involved in many cellular processes^21,22^ and were recently identified as key initiators of post-ischemic inflammation.^23^ hPrx5 therefore emerges as a target for the development of treatments against strokes, and has been the object of screening campaigns. Notably, we previously screened hPrx5 by NMR and discovered fragment molecules from the catechol family as active-site ligands and potential inhibitors.^24^ Later on, we described their binding/unbinding to hPrx5 using unbiased MD and FM, achieving a free-energy prediction that matched quantitatively the low affinity determined by high-accuracy solution NMR.^16^ Further, we correlated the binding affinities of catechol, 4-methylcatechol (4MEC) and 4-tert-butylcatechol (4TBC) to hPrx5 to a reversible inhibition mechanism using steady-state kinetics, demonstrating a partial mixed-type non-competitive mechanism, in agreement with FM simulations in the presence of the substrate.^25^

In these previous studies, the simulations indicated that the rather rigid ligands (CAT, 4MEC, 4TBC) could adopt several bound state conformations (three conformations bound to the catalytic cysteine via one or two hydrogen bonds, as well as a conformation bound to the Glu179 side-chain in the active site cavity), and that the fluctuation of the activesite geometry was affected during the binding periods.^25^ Whereas 4MEC binding caused, on average, a reduction in the volume of the active site,^16^ 4TBC caused a global expansion during the binding periods as seen in Fig S1 around 300 ns of simulation. From these observations, we hypothesized that the introduction of an additional degree of freedom in the substituent of catechol could lead to an improved affinity and inhibition. Indeed, we reasoned that it would help the ligand better adapt to the active-site dynamics, and hence reduce the conformational dispersion during the binding periods by stabilizing a binding mode relative to the others. Through simulations and experiments, we show here that this is in fact the case: an additional degree of freedom in the ligand structure — lengthening the aliphatic tail by one carbon atom to obtain the corresponding 4-neo-pentylcatechol (4NPC) — improves the inhibition of hPrx5 by several orders of magnitude.

## Results

### Funnel Metadynamics predicts improved affinity of 4NPC for hPrx5

To test our hypothesis, we performed careful free energy simulations with the Funnel Meta-dynamics (FM) method to assess whether 4NPC does exhibit an improved affinity for hPrx5. In a 1 µs FM simulation with 4NPC, we observed 6 reversible binding events. These observed recrossings, along with the stabilization of the free energy estimation in the second half of the simulation, supports the proper convergence of this challenging calculation. This allowed us to evaluate the absolute binding free energy 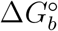 (from the free-solvated state to the bound form) of this complex at the convergence of the calculation.^15^

The estimated 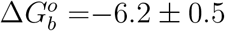 kcal mol^−1^ predicts an improved affinity in comparison with all previous catechol derivatives (Table 1).^16,25^ Moreover, we note that the binding events are of various durations, with the longest lasting ≈ 100 ns (see S1 and S2), *i*.*e*.,, longer than for previous studied ligands.^16,25^ This observation suggests that 4NPTC may also exhibit an increased residence time in the active site of hPrx5, which is a desirable property for an inhibitor. Thus, FM provides us with a quantitative estimation of the affinity of the complex, which encouragingly predicts that the proposed derivative should indeed exhibit improved affinity towards the target. Next, to unravel the atomistic determinants of improved binding, we analyze the FM trajectory and compare it to previous simulations of other hPrx5 binders.

**Table 1:** Comparison of the predicted and the experimentally determined affinities and inhibitions of catechol derivatives against hPrx5.

|  | Catechol | 4TBC | 4NPC |
| --- | --- | --- | --- |
| Affinities |  |  |  |
| $\Delta G_b^\circ$ (kcal mol<sup>-1</sup>)<sup>c</sup> | $-3.0 \pm 0.2^a$ | $-5.5 \pm 0.9^b$ | $-6.2 \pm 0.5$ |
| $K_D$ (μM) - FM<sup>c</sup> | $6900 \pm 2100^a$ | $110 \pm 90^b$ | $60 \pm 30$ |
| $K_D$ (μM) - NMR<sup>c</sup> | $4500 \pm 600^a$ | $190 \pm 30^b$ | $20 \pm 10$ |
| Inhibitions |  |  |  |
| IC50 (μM) | $3730 \pm 850$ | $250 \pm 60$ | $40 \pm 5$ |
| $K_M$ (μM) | $40.7 \pm 4.0^b$ | $36.9 \pm 4.4^b$ | $26 \pm 2$ |
| $K_i$ (μM) | $1810 \pm 460^b$ | $90 \pm 40^b$ | $19 \pm 5$ |
| $K'_i$ (μM) | $4040 \pm 1130^b$ | $180 \pm 100^b$ | $80 \pm 40$ |
| $k_{cat}$ (s<sup>-1</sup>) | $3.08 \pm 0.08^b$ | $3.05 \pm 0.10^b$ | $2.15 \pm 0.05$ |
| $k'_{cat}$ (s<sup>-1</sup>) | $\sim 0^b$ | $2.35 \pm 0.18^b$ | $0-0.67^d$ |

<sup>a</sup> Data from a previous report.;<sup>16</sup> <sup>b</sup> Data from a previous report.;<sup>25</sup> <sup>c</sup> In FM, $\Delta G_b^\circ$ and $K_D$ were calculated by standard deviation, reweighted over the simulation time (last 100 ns over 500 ns for catechol and 4TBC, and last 400 ns over 1000 ns for 4NPC). In NMR, $K_D$ was calculated from CSPs.; <sup>d</sup> For $k'_{cat}$ we determined an upper limit of $0.67 \text{ s}^{-1}$ .

### Ligand dynamics within the active site

We analyzed the trajectory to get insight into the conformational dynamics of this new ligand in the active site, and how it can be related to affinity improvement. For this purpose, we first projected the 4NPC FM trajectory onto three collective variables (CVs): CV1, the ligand-active site distance, the one biased in the FM calculation; and CV2 and CV3 that are designed to capture the conformational state of the ligand (CV2) and its orientation relative to the active site (CV3); see Methods section for more information on CVs. Thus, our protocol provides direct insight into how binding to the protein shapes the conforma-tional landscape of the ligand. Using the reweighting procedure described in reference^26^ as implemented in PLUMED,^27^ we mapped the free energy as a function of CV1 and CV3. The analysis of this free energy landscape (Fig. 2) reveals three local free energy minima in the bound state. Structural clustering shows that these correspond to four major binding conformations A, B, C and D. Three of them (A,B,C) interact directly through H-bonding with the catalytic thiolate of Cys47 while the neo-pentyl group makes Van der Waals lateral contacts to the hPrx5 active site region including Leu116, Ile119 and Phe120 (2). Remarkably, the stabilisation of these bound conformation via lateral contacts with the hydrophobic helix 116-120 is now possible thanks to the added degree of freedom. The fourth cluster (D) retains these lateral interactions while making stabilizing polar interactions with Glu179 at the surface of hPrx5, as previously detected for 4TBC,^25^ and is 1 kcal mol^−1^ higher in free energy. Overall, in comparison with previous catechol fragments that were exhibiting several dispersed bound conformations (6 for 4TBC), only four different conformers are detected for the 4NPC, all within 1 kcal mol^−1^ of one another and separated by low (≈2 kcal mol^−1^) barriers, suggesting that they exchange rapidly and all contribute to the bound state. Moreover, we observe that in comparison with previous ligands, 4NPC does not induce significant changes into the active site fluctuations, as depicted in the lower panels of Fig. S1. This suggests that 4NPC exhibits better flexibility and adaptation to the active site geometry, which contributes to the increased duration of binding periods as well as additional stabilizing contacts observed in clusters A,B and C. These improvements should lead to an overall, experimentally detectable better affinity and inhibition of the protein by the 4NPC.

**Figure 1:**
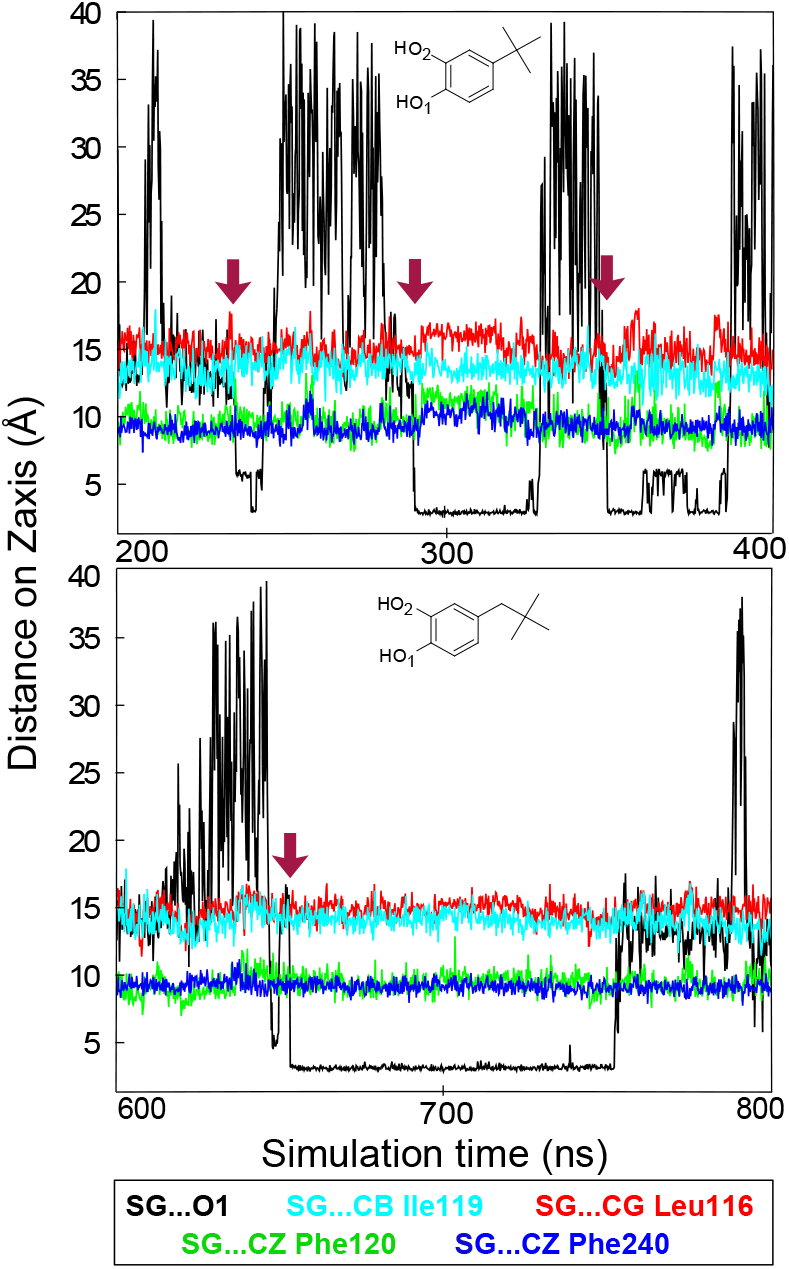
Comparison of the interatomic distances as a function of the simulation time for 4TBC (upper pannel) and 4NPC (lower pannel) ligands. Shown are windows of 200 ns from simulations, with different binding periods. Protein-ligand distance is plotted in black, measured between the sulphur atom of the Cys47 and the O1 atom of the ligand. Binding events are indicated with red arrows. The following interatomic distances within Prx5 active site are plotted in colors : SG Cys 47 to CG Ile 119 in cyan, SG Cys 47 to CG Leu 116 in red, SG Cys 47 to CZ Phe120 in green and SG Cys 47 to CZ Phe240 in blue.

**Figure 2:**
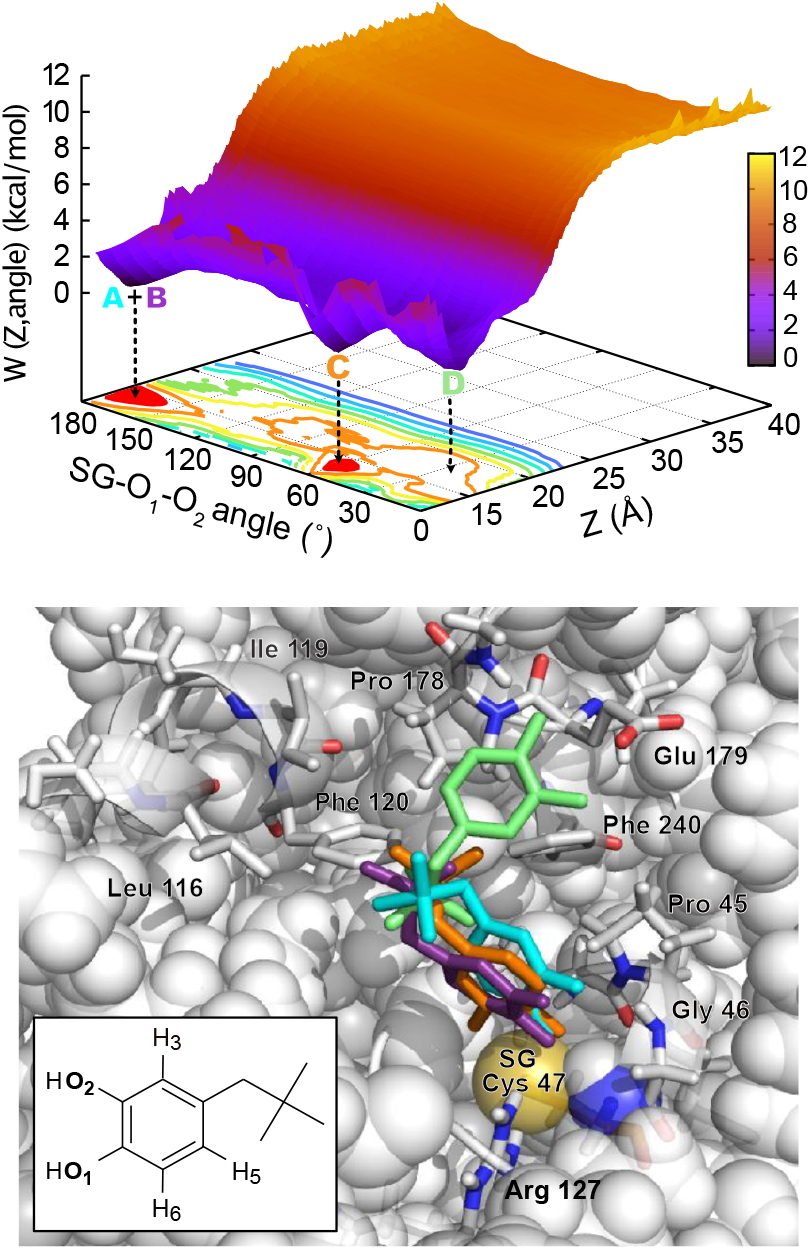
Free-energy surface (FES) and geometry of the 4NPC-hPrx5 interaction. Upper panel: FES as a function of two collective variables used in FM calculations. CV1: projection on the *Z* axis of the distance between Cys47 SG and the center of mass of the ligand heavy atoms (Å) and CV3: SG-O1-O2 angle (deg) between sulphur atom of the catalytic cysteine and oxygen atoms of the diol ligand. Iso-energy contours are drawn using 1 kcal mol^−1^ interval. Lower panel: visualisation of the four bound conformations obtained after geometrical clustering of the FM trajectory with a 2 Å cutoff, representative of the three free-energy basins A+B, C and D. Cys47 is shown as a yellow Van der Waals sphere and clusters conformations are colored as follows: A: blue, B: purple, C: orange, D: green.

### NMR assay confirms the improved affinity of 4NPC for hPrx5

To test the FM results experimentally, we measured the affinity of 4NPC for hPrx5 by solution NMR assays (3). The ^1^ H-^15^ N chemical shift perturbations (CSP) observed on ^1^ H-^15^ N-HSQC spectra in the presence of 4NPC were successfully fitted to a simple binding model (see Methods), allowing us to estimate the binding affinity. We found an NMR-based average dissociation constant *K*_*D*_ = 20 *±* 10 µM. This value falls on the lower range of the statistical fluctuations of the FM prediction. Furthermore, the CSPs exhibited a greater magnitude compared to those observed with other catechol fragments (Fig. S3), confirming less dispersed bound conformations. A comparison of the CSPs induced in the case of 4TBC and 4NPC reveals that the enhancement in affinity is accompanied by a greater degree of disruption of the Leu116-Phe120 hydrophobic helix, as well as the residues located in the vicinity of Cys47. This increase in the CSPs reflects the enhanced stability of the bound form of the PL complex. Finally, an NMR saturation transfer difference (STD) epitope mapping (Fig. 3) reveals that protons H5 and H6 exhibit a much larger degree of saturation transfer (respectively 90% and 100%) than proton H3, suggesting that H5 and H6 are more involved into the protein-ligand interface whereas H3 is more solvent-accessible. These characteristics are remarkably consistent with the ligand poses observed in clusters A and B (Fig 2), which were predicted as dominant from the simulations. Therefore, NMR measurements are consistent with both the simulation-based affinity prediction and the ligand/receptor conformational landscape observed in the simulation.

**Figure 3:**
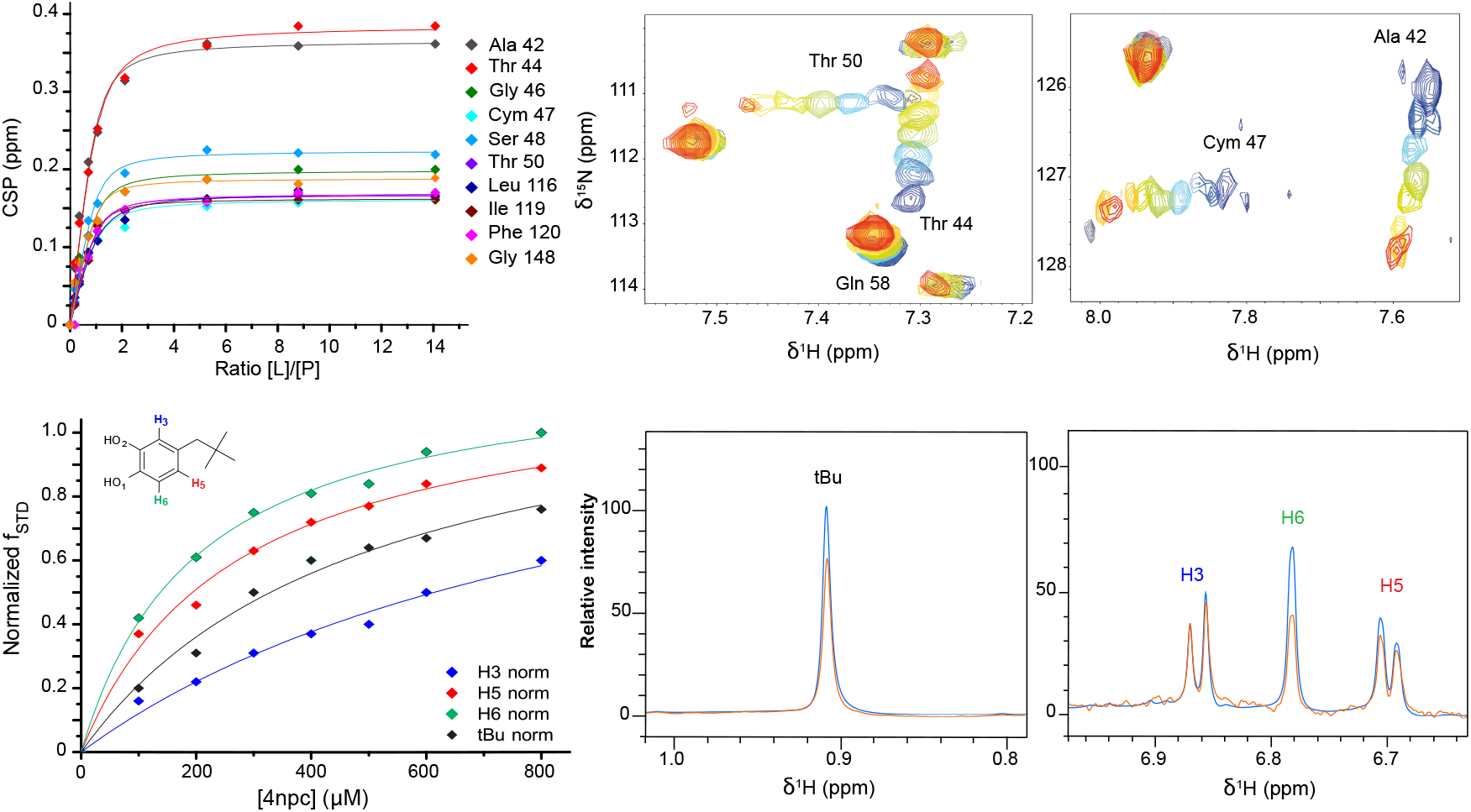
Affinity assay of 4NPC for hPrx5 using NMR. Upper panels: plot of the NMR CSP obtained as a function of ligand-to-protein concentration ratio, and two partial overlays of ^1^ H-^15^ N HSQC spectra of hPrx5 with various concentrations of 4NPC in PBS pH=7.4 at 301 K, from blue (hPrx5 only) to red (saturation of the binding site by 4NPC). Lower panels: plot of the STD factor against ligand concentration for aromatic protons and tert-butyl group, and spectral overlays of the 1D spectra (blue) and the STD spectra (orange) obtained for [4NPC] = 0.8 mM.

### Inhibition mechanism and enzymology

Next, we set out to find out whether the increased binding affinity actually translates into an improved inhibitory effect as compared to previously characterized catechol derivatives. For this purpose, we conducted experiments to evaluate the reversible mechanism of inhibition for 4NPC, and compared it to other catechol fragments.^25^ First, our enzymatic assay yielded an IC50 of 40 ± 5 µM for 4NPC, considerably lower than for catechol (3730 ± 850 µM) and the tert-butyl derivative (250±60 µM), see Table 1. Then, we studied the mechanism of inhibition using steady-state kinetics analyzed with non-linear regression models.^28,29^ Reaction rates were fitted against the concentration of H_2_O_2_ for each concentration of 4NPC and found to fit best with a partial mixed non competitive inhibition model. In this model, 4NPC can either bind to the reduced-free enzyme and to the Enzyme-Substrate (ES) complex (Fig. S5) with two different affinities (*K*_*i*_ and 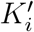). Based on the values obtained for inhibition constants (*K*_*i*_ and 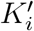) and catalytic rates (*k*_*cat*_ and 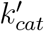) reported in Table 1, 4NPC binds preferentially to the free reduced enzyme 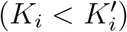 and the ternary ESI complex is inhibited 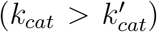. The inhibition constants (*K*_*i*_) indicate that 4NPC inhibits the free reduced hPrx5 100 times better than catechol, and 5 times better than 4TBC, demonstrating that we have obtained an improved inhibitor through rational modification.

## Discussion and Conclusion

Protein-ligand affinity results from a complex interplay between strong interactions and conformational plasticity. Small, rigid ligands tend to favor strong binding by reducing entropic penalty, but the affinity gain can be offset if the ligand is too rigid to interact optimally with the binding site. Here, we improved the activity of an inhibitor by tuning flexibility to promote conformational adaptation to the target active site (Fig. 4). The introduction of one degree of freedom in 4NPC results in a better adaptation to the active site through the formation of additional intermolecular interactions besides the binding to the catalytic cysteine. Interestingly, although the binding affinity of 4NPC for hPrx5 (both predicted and measured) is only marginally better than that of 4TBC,^25^ the inhibitory effect (*e*.*g*., the IC50, Table 1) is markedly improved. Based on our simulation observations, we hypothesize that this may be due to an increased residence time of 4NPC thanks to stabilization of the main binding mode. This is consistent with recent evidence that the residence time is a better activity predictor than the dissociation constant.^30^

**Figure 4:**
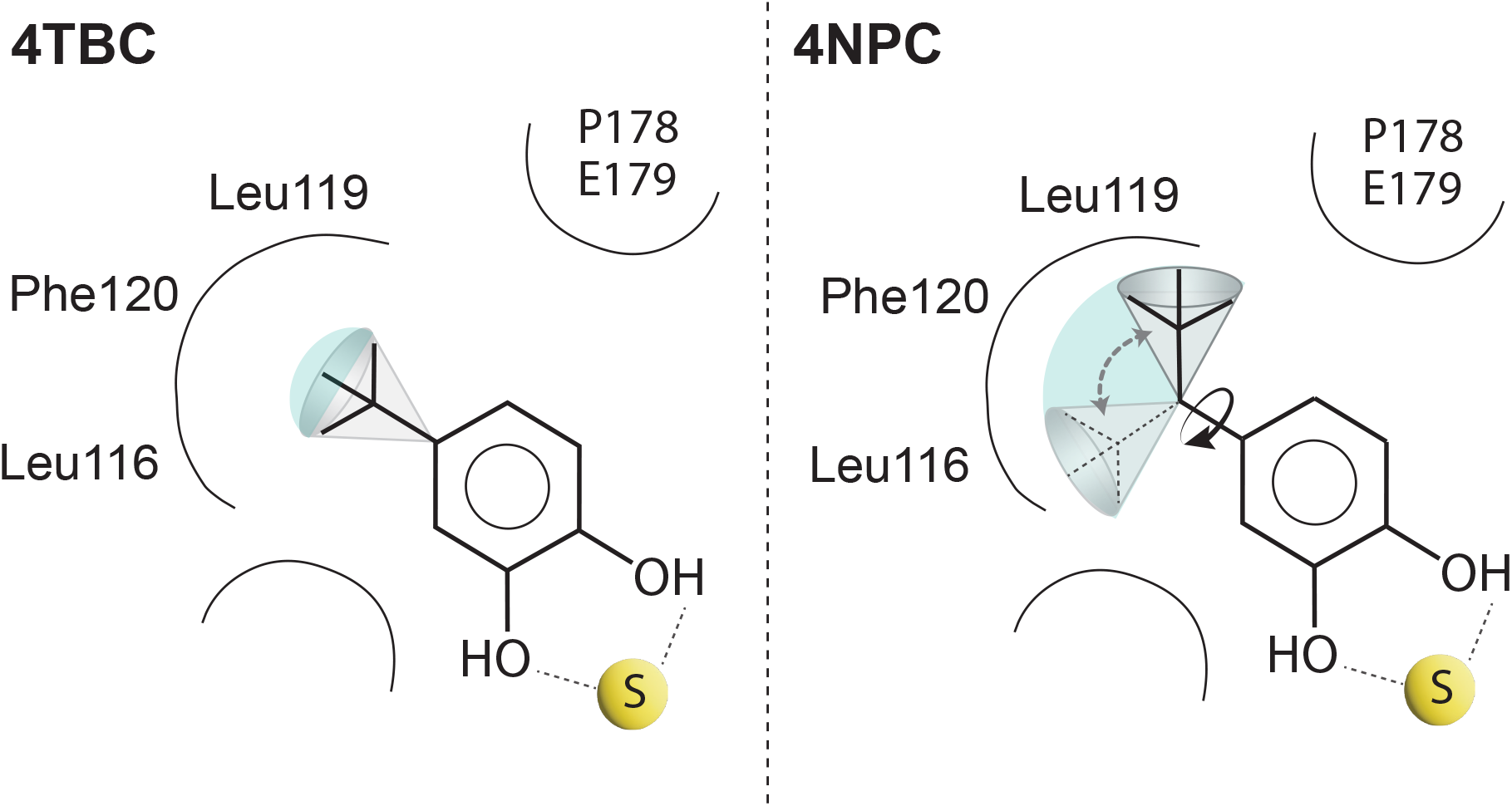
Chemical modification to tert-butyl-catechol led to improved spatial adaptation.

We introduced what is arguably the simplest, minimum structural modification to that effect. This leaves a lot of unexplored chemical space to evolve toward a drug-like compound with further improved, nanomolar inhibitory properties against hPrx5. We expect the insights into protein-ligand interactions afforded by FM to be precious for this next optimization step.

## Methods

### Protein expression and purification

Recombinant human Prx5, yeast Trx, and yeast TrxR were inserted in the pQE-30 expression vector (Qiagen) with an N-terminal fusion with hexahistidine (6xHis) tag. The expression and purification of the proteins were adapted from Declercq et al.^31^ The plasmids kindly provided by Dr. Bernard Knoop’s lab (Université Catholique de Louvain, Belgium) were transformed into *Escherichia coli* strain M15 (pRep4) and grown in LB medium at 37 ^°^C. To produce ^15^ N labeled human hPrx5, cells were expressed in M9 minimal medium (6 g/L Na_2_HPO_4_, 3 g/L KH_2_PO_4_, 0.5 g/L NaCl) supplemented with 6 µg thiamine, 1 mM MgSO_4_, 1 mM CaCl_2_, 10 mL/L trace metal solution, 4 g/L glucose, two resistance antibiotics (kanamycin and ampicillin 50 µg mL^−1^) and containing 1 g/L ^15^ NH_4_Cl as the sole nitrogen source. The bacterial cultures were induced for an O.D. of 0.6 with 1 mM of isopropyl -D-1-thiogalactopyranoside (IPTG) for 4 to 5 hours and were then centrifuged for 20 min at 3000 rpm. The resulting bacterial cell pellets were lysed by sonication using a solution containing 10 mM imidazole, 50 mM phosphate and 300 mM NaCl pH 8 (lysis buffer). The cell lysate was then centrifuged for 45 min at 12000 rpm and the supernatant loaded into a Ni^2+^-NTA column (Qiagen). The column was washed with lysis buffer and the protein was eluted with 250 mM imidazole, 50 mM phosphate, 300 mM NaCl pH 8. The eluted protein was pooled and dialyzed overnight against PBS pH 7.4 at 4 ^°^C, then analyzed by SDS-PAGE. Its final concentration was quantified by UV-Vis spectrum (A280: Trx (*ε* = 10095 M^-1^ cm^-1^), TrxR (*ε* = 24660 M^-^1 cm^-1^), and human Prx5 (*ε* = 5625 M^-1^ cm^-1^) with *ε* calculated from Expasy website (expasy.org/protparam).

### Molecular Dynamics Simulations

MD systems were prepared, minimized, and equilibrated as described previously^16^ from the high-resolution crystal structure of homodimeric hPrx5 (PDB entry: 3MNG).^32,33^ Crystallographic water molecules were retained and added to the protein coordinates. Hydrogen peroxide and ligands were positioned manually near one of the two active sites of the homodimer, and the catalytic cysteine (Cys47) was modeled as a thiolate. Finally, Na^+^ and Cl^−^ were added to match 150 mM sodium chloride aqueous solution, close to the experimental conditions used, and to neutralize the total charge of the system. We assumed that all residues were in their standard protonation state at pH 7.4 (physiological conditions).

Simulations were performed with the AMBER99SB-ILDN force field^34–36^ for the protein, and the TIP3P water model for the explicit solvent.^37^ We used the General Amber Force Field (GAFF)^38^ bonded parameters and charges for the ligands, which we prepared using the Antechamber program suite.^39^ The system was energy-minimized and equilibrated under constant pressure and temperature (NPT) conditions at 1 atm and 300 K using ACEMD as described in reference,^6^ and the PLUMED plugin was used to carry out funnel-metadynamics calculations.^27^ Since the funnel external potential was fixed in space, diffusion of the whole protein was canceled by harmonically restraining the positions of 3 atoms chosen far from the interface of the homodimer or from the active site, namely the C atoms of G6, G31, and K65 of chain A, with a force constant of 20 kcal*/*mol*/*Å^2^ in ACEMD. The geometric parameters of the funnel are as follows: *h*_*cyl*_ = 17 Å; *R*_*cyl*_ = 1 Å; *h*_*cone*_ = 18 Å; *α* = 1.1 rad; *h*_*funnel*_ = *h*_*cone*_ + *h*_*cyl*_. Additional details are provided in Troussicot et al. 2015.^16^

We evaluated the free energy profile along the distance of the ligand to the center of the active site by well-tempered metadynamics in the presence of the funnel potential. Three collective variables (CV) were defined : the projection on the *Z*-axis of the distance between Cys47 SG and the center of mass of the ligand heavy atoms (CV1), the distance from the funnel Z-axis of the ligand center of mass (CV2) and the protein-ligand torsion angle Cys47 SG…O1-C1-C2 (ligand). The bias was applied on CV1, and the values of all CVs were collected along the simulations to evaluate the conformational space explored by the system. A Gaussian width of 0.35 Å and deposition rate of 0.25 kcal mol^−1^ ps^−1^ (1 kcal = 4.18 kJ) were initially used and gradually adapted using well-tempered metadynamics with a bias factor *γ* =12 (that is, Δ*T* =3600 K).^7^ At convergence, the binding free energy was recovered from the Potential Mean Force after correcting for the entropic penalty of the funnel following the standard procedure described in.^15^ Convergence was assessed by monitoring the time-evolution of the Δ*G* estimate. Specifically, we considered the free energy estimate to be converged when residual fluctuations lower than 1 kcal mol^−1^ were reached. Free energy surfaces along CVs different than CV1 (*i*.*e*.,, CVs not directly biased by well-tempered metadynamics) were evaluated following the reweighting procedure described in.^26^ The dissociation constants *K*_*D*_ were computed as 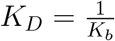 with *K*_*b*_ = exp(−Δ*G*^°^*/RT*) the binding constant.

### Chemical Synthesis of the 4-neopentylbenzene-1,2-diol

The 4-neopentylbenzene-1,2-diol **4** was synthesized from 1,2-dimethoxybenzene **1** according to the scheme depicted in Fig. 5. The Friedel-Crafts acylation was performed in an ionic liquid^40^ with Bi(OTf)_3_ and the resulting t-butylphenylketone **2** was reduced to the corresponding alkane **3** under green conditions (cyclopentyl methyl ether - CPME) and Pd/C 5% as catalyst.^41^ The final removal of the methyl ethers with BBr_3_ yielded pure 4NPC product **4** as a white crystalline powder. The details of the synthesis steps are available in SI (Supplementary Text 1), as well as the mass spectrometry and NMR analysis of the different intermediates, and the corresponding ^1^ H and ^13^ C NMR spectra (Figs S6, S7 and S8).

**Figure 5:**
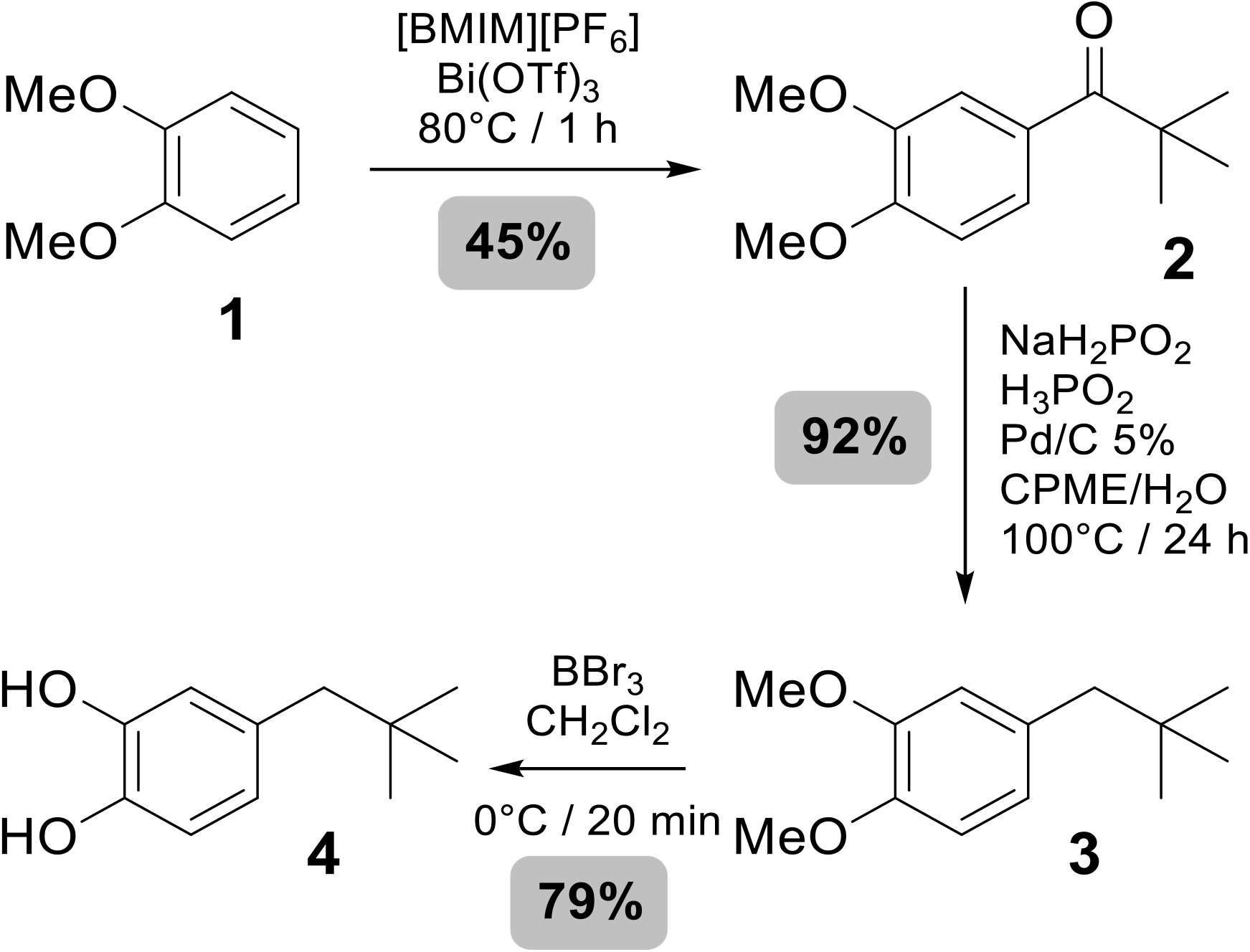
Scheme: Chemical synthesis of 4-neopentylbenzene-1,2-diol and the yields obtained for the different steps of the synthesis.

### Solution NMR experiments

HSQC experiments were monitored at 28 ^°^C with a Varian Inova 600 MHz NMR spectrometer equipped with a 5 mm standard triple resonance (^1^ H/^13^ C/^15^ N) inverse probe with a z-axis field gradient, whereas STD standard 1D and STD experiments were performed on the same Varian Inova 600 MHz NMR spectrometer further equipped with a 5mm PFG triple resonance HCN cold probe at 28 ^°^C. NMR pulse sequences used for these experiments were taken from the Agilent Biopack ensemble of experimental schemes within Agilent VnmrJ 4.2 software. For HSQC experiments, a Watergate water suppression HSQC sequence and for STD experiment a Saturation Transfer 1D sequence were chosen. The binding interaction of 4NPC to Prx5 was characterized through a series of two-dimensional ^1^ H-^15^ N-HSQC NMR experiments. NMR samples contained 200 µM of the reduced ^15^ N-labeled protein, kept in its reduced form by addition of 2 mM of tris(2-carboxyéthyl)phosphine (TCEP), in phosphate buffer saline (PBS) pH 7.4. An increasing concentration of 4NPC previously dissolved in dimethyl sulfoxide (DMSO) was added until binding saturation was reached. DMSO’s influence on chemical shift was tested until 8%. All spectra were processed using NMRPipe/NMDraw package^42^ and ^1^ H-^15^ N chemical shift perturbations (CSPs) were assigned using NMRViewJ software.^43^ The combined amide CSPs Δδ was calculated accordingly to the following expression:

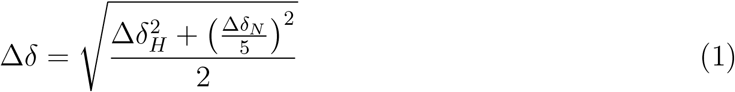

Upon characterizing the amide CSP Δδ to the specific amino acid residues involved in the binding interactions, a binding dissociation constant (K_D_) was extrapolated from the curve plotting the CSP values versus the concentration ratio [Ligand]/[Prx5] as previously described by Lange et al. 2011.^44^ The binding of 4NPC to Prx5 was also investigated using 1H Saturation Transfer Difference (STD) NMR spectroscopy. NMR samples were prepared in PBS (pH 7.4) with 10 µM Prx5 in presence of TCEP (500 µM) to keep it in reduced form, and 4NPC was added from 100 µM to 1 mM. 1D and Saturation Transfer Difference (STD) experiments were acquired using identical conditions, and all STD spectra were recorded with the same parameters (saturation period = 2.0 s, made of 40 Gaussian pulses of 50 ms). The number of scans was set to 512 for 1D spectra and 1024 scans for STD, and the selective irradiation of the protein was performed with a decoupler offset of -3006 Hz downfield from the carrier frequency (-0.23 ppm), and off-resonance control was performed at 15000 Hz upfield. STD signals were then measured for aromatic protons and tert-butyl group, and the STD factor was calculated and then normalized by setting the largest value, following the formula:

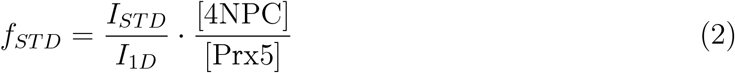

where *f*_*STD*_ is the STD factor, *I*_*STD*_ is the integral of a given STD signal, *I*_1*D*_ is the integral of the normal spectrum of the same signal, and [4NPC]/[Prx5] is the ligand to protein concentration ratio.

### Half Maximal Inhibitory Concentration Assay (IC50)

The inhibitory concentration assay of Prx5 was carried out as described by Chow et al. 2016.^25^ The inhibition activity of Prx5 with 4NPC was indirectly measured by the coupling reaction of thioredoxin reductase (TrxR) oxidizing Nicotinamide adenine dinucleotide phosphate (NADP) monitored with the absorbance of NADPH at 340 nm (A_340_) (see Supporting Information S6). Within a total of 1000 µL, the reaction solution consists of 4NPC at 0 µM to 1500 µM range, 15 µM of yeast thioredoxin (Trx), 2 µM of yeast TrxR, 0.15 µM of human Prx5, 200 µM of NADPH and 25 µM of H_2_O_2_. The reaction was carried out by a mixture of all three proteins diluted in PBS containing 1mM of ethylenediamine tetraacetic acid (EDTA) pH 7.0 and the successive addition of 4NPC and NADPH. Lastly, H_2_O_2_ was added to the mixture and mixed well to initiate the inhibition activity of Prx5 at room temperature. The reaction was monitored at A_340_ for 200 seconds on a Jasco UV-Visible spectrometer, and the initial rate of the reaction was determined from linear portion of the curve and is expressed in µmol min^−1^ mg^−1^ of Prx5.

### Inhibition mechanism assay

Assays to determine the inhibition mechanism of 4NPC for human Prx5 was adapted from Chow et al. 2016.^25^ Four different concentrations of 4NPC were used, taken above and below the value of IC50 (40 ± 5 µM). Reaction was carried out by mixing together Prx5 (0.15 µM), yTrx (15 µM), yTrxR (2 µM), the catechol derivative, and NADPH (200 µM). The reaction was started with the addition of H_2_O_2_ (range 0 µM to 500 µM) in the reaction mixture. The reaction was monitored with the same device and parameters than for the IC50 determination, and the rate expressed in µmol min^−1^ L^−1^.

## Supporting information

Supplementary Informations

## Supporting Information

Are presented in the Supporting Information: supplementary figures showing the trajectory analysis of the reduced Prx5 in presence of 4NPC, the active site fluctuations observed during the simulations of Prx with ligands 4TBC and 4NPC, the histogram of the chemical shift perturbations (CSPs) induced by the biding of the different catechol derivatives with Prx5, the IC50 determination for the 4NPC ligand, the Partial-Mixed Type Non Competitive Inhibition Model, the analysis of the inhibition mechanism assays realised on the 4NPC, and the ^1^ H and ^13^ C NMR spectra obtained for the synthesized compounds (intermediates and final product). Supplementary Text 1 describes the chemical syntheses.

## Abbreviations

CAT: catechol
4MEC: 4-methylcatechol
4TBC: 4-tertbutylcatechol
4NPC: 4-neopentylcatechol
Prx5: Human Peroxiredoxin 5

## Acknowledgements

We acknowledge la Région Rhônes Alpes for the grant no. 13-0118962-01 allocated to L.T. and J-M.L, and the Institut de Chimie de Lyon (2016) for additional funding. We are grateful to Prof. B. Knoops and A. Clippe from University of Louvain-la-Neuve, Belgium, for the generous gifts of biological materials to express and purify the recombinant proteins. We thank Acellera Company for providing of a 6-month ACEMD multi-GPU software license and for the gift of a Nvidia Tesla K20 supercomputing unit. Finally, we thank Dr. Maggy Hologne for useful discussions.

