## Supplementary Informations for "Metadynamics Simulations meet Ligand Design for the Reversible Inhibition of Human Peroxiredoxin 5"

### Supplementary Text 1 : Details of the chemical synthesis of the 4NPC ligand

Chemicals were obtained from commercial sources and used without further purification. Reactions were performed under an argon atmosphere. Thin-layer chromatography (TLC) was carried out on aluminum sheets coated with silica gel 60 F254 (Merck). TLC plates were inspected by UV light (  = 254 nm). Silica gel column chromatography was performed with silica gel Si 60 (40 µm to 63 µm). NMR spectra given as supplementary information were recorded at 303 K using a Bruker Avance III HD 400 MHz spectrometer. Chemical shifts ($\delta$ in ppm) are referenced relative to deuterated solvent residual peaks. The following abbreviations are used to explain the observed multiplicities: s, singlet; d, doublet; t, triplet; q, quadruplet; m, multiplet. Complete signal assignments were based on 1D and 2D NMR (COSY and HSQC correlations). High-resolution (HR-ESI-QToF) mass spectra were recorded using a Bruker MicroToF-Q II XL spectrometer.

**1-(3,4-Dimethoxyphenyl)-2,2-dimethylpropan-1-one(2)**: 1,2-Dimethoxybenzene 1 (10 mmol, 1.27mL), $Bi(OTf)_3$ (0.5 mmol, 328 mg) and pivaloyl chloride (10 mmol, 1.23 mL) were stirred at 80C in [BMIM][PF6] (5g) during 1 h until disappearance of the starting material (TLC monitoring: eluent $CH_2Cl_2$/petroleum ether, 3:2). After cooling, the reaction mixture was extracted with diethylether (4×50 mL) using sonication to help separating the organic phase from the ionic liquid. The ether layers were combined then washed with brine (4×30 mL), dried ($Na_2SO_4$) and concentrated. The crude residue was purified by flash chromatography ($CH_2Cl_2$/petroleum ether, 1:1 then 4:1) to give 1-(3,4-dimethoxyphenyl)-2,2-dimethylpropan-1-one 2 (1g, 45%) as a yellow oil. $R_f$= 0.21 ($CH_2Cl_2$/petroleum ether, 3:2). HR-ESI-MS: m/z calculated for $C_{13}H_{18}NaO_3$ $[M+Na]^+$ 245.1148, found 245.1145.

**4-(2,2-Dimethylpropyl)-1,2-dimethoxybenzene (3)**: In a sealed vial was introduced ketone 2 (2.34 mmol, 520 mg) dissolved in cyclopentylmethyl ether (2.5 mL) and distilled water (5 mL). Pd/C 10% (0.234 mmol, 249 mg), sodium hypophosphite monohydrate (7.02 mmol, 744 mg) and hypophosphorous acid (1.17 mmol, 140 mg, 50% wt) were added. The resulting mixture was heated at 100C under inert atmosphere (Argon) during 24 h. The reaction was diluted with $CH_2Cl_2$ (30 mL) and water (40 mL). The aqueous layer was then extracted with $CH_2Cl_2$ (4×75mL). The organic layers were combined, dried ($Na_2SO_4$), filtered over a pad of celite then washed with $CH_2Cl_2$ (100 mL). The solvent was removed on a rotary evaporator to provide 4-(2,2-dimethylpropyl)-1,2-dimethoxybenzene 3 (448 mg, 92%) as a yellow oil-without purification. The product crystallized in the fridge (4C) after a few hours as a pale yellow solid. $R_f$= 0.77 ($CH_2Cl_2$/petroleum ether, 7:3). M.p.= 36-37C. HR-ESI-MS: m/z calculated for $C_{13}H_{20}NaO_2$ $[M+Na]^+$ 231.1356, found 231.1351.

**4-(2,2-Dimethylpropyl)benzene-1,2-diol (4)**: A solution of $BBr_3$ (4.5 mmol, 4.5 mL, 1M in $CH_2Cl_2$) was added under inert atmosphere at 0C to a solution of compound 3 (1.4 mmol, 290 mg) in $CH_2Cl_2$ (15 mL).The dark red/brown reaction was stirred at 0C during 20 minutes and then quenched by adding a saturated aqueous solution of $NaHCO_3$ (15 mL) and stirring was continued for an additional 2.5 h. After dilution in $CH_2Cl_2$ (60 mL), the organic solution was extracted with 10% aqueous $Na_2S_2O_3$ (50 mL) to remove bromine, and washed with ice-cold water (30 mL), dried ($Na_2SO_4$) and filtered. The solvent was removed and the crude product was purified by a silica gel flash chromatography ($CH_2Cl_2$/MeOH, 98:2) affording the 4-(2,2-dimethylpropyl)benzene-1,2-diol 4 (198 mg, 79%) as a white shiny powder. $R_f$= 0.17 ($CH_2Cl_2$). M.p.= 124-125C. HR-ESI-MS: m/z calculated for $C_{11}H_{15}O_2$ $[M-H]^-$ 179.1078, found 179.1071.

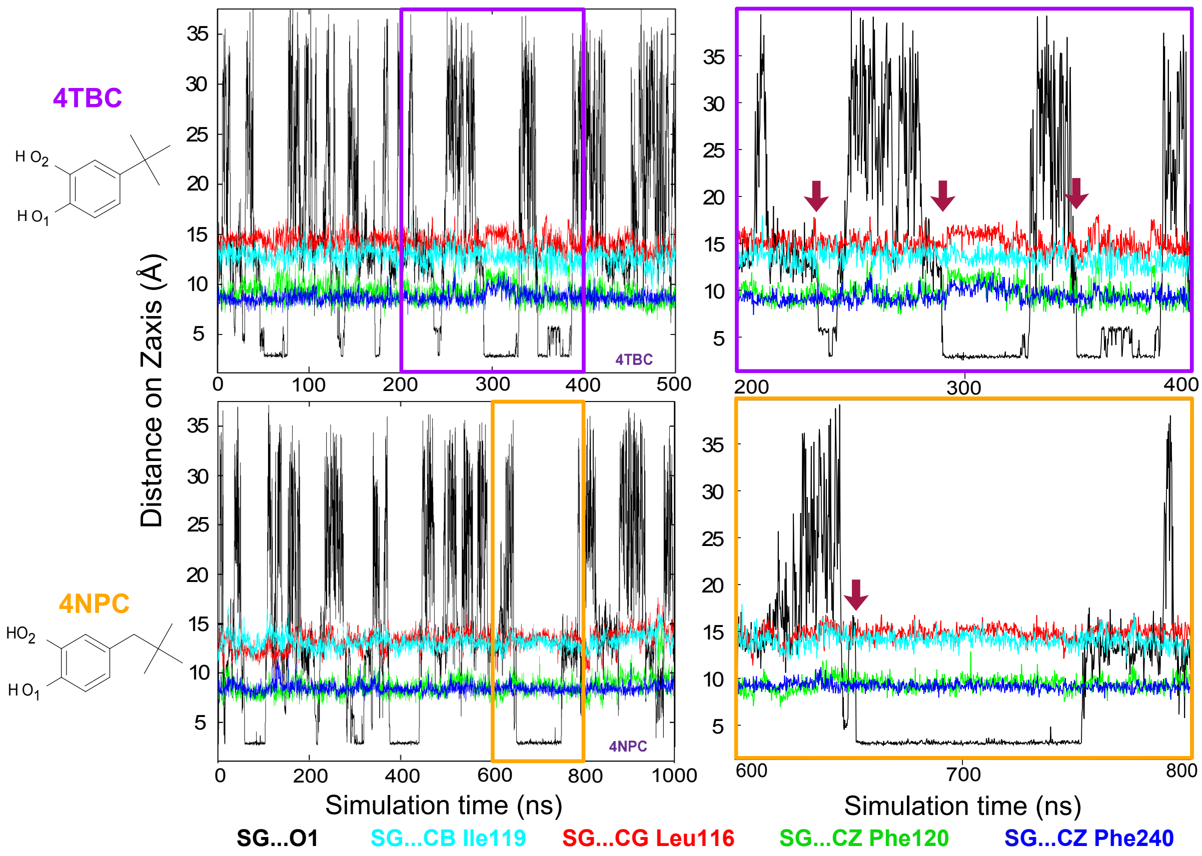


Figure S1: Comparison of the interatomic distances as a function of FM simulation time for 4TBC (top panels) and 4NPC (lower panels) ligands. Protein (Cys47)-ligand distance (black trace), measured between sulphur atom of the cystein (SG) and O1 atom of the ligand. Binding events are indicated with red arrows. Interatomic distances within hPrx5 active site are plotted in cyan (SG...CG Leu 116), red (SG...CB Ile 119), green (SG...CZ Phe 120) and blue (SG...CZ Phe 240) against the entire simulation time (500 ns for 4TBC and 1 µs for 4NPC) on the left panels. On the right panels, a zoom of a 200 ns period is depicted to compare the duration of the binding period and its effect on the interatomic distances.

.

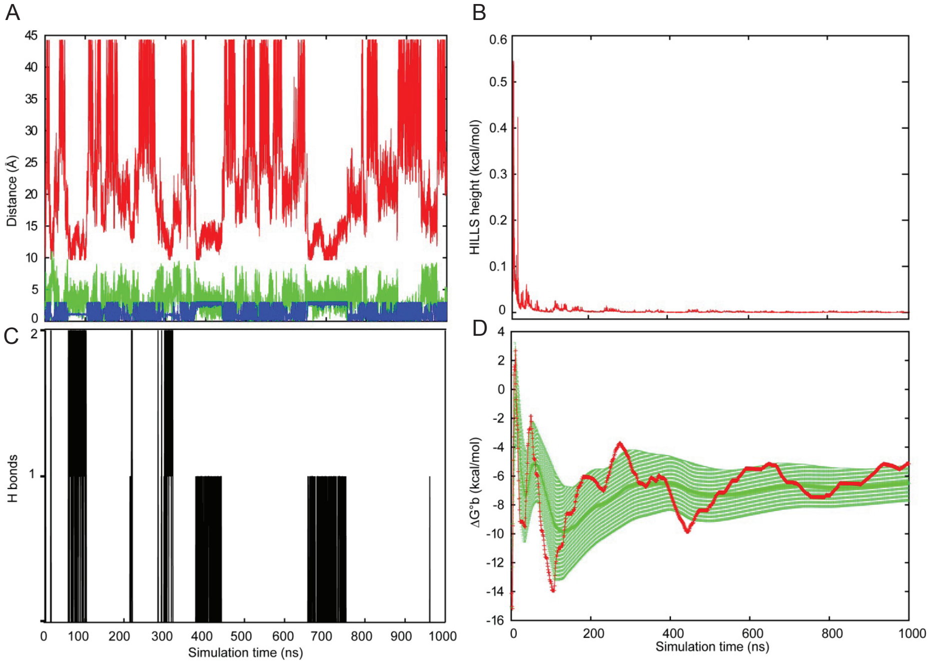


Figure S2: MD informations. A) representation of the values of the 3 collective variables used in the MD simulation as a function of the simulation time. Red trace (CV1): projection on the Z axis of the distance between Cys47 SG and the center of mass of the ligand heavy atoms (Å). Green trace (CV2): distance of the ligand's center of mass from the Z axis (Å). Blue trace (CV3): Cys47 SG...O1...O2 angle between catalytic cysteine and oxygen atoms of the ligand (rad). B) Hills potential deposition in kcal/mol along the simulation time. C) Counting of the hydrogen bonds formation between Cys47 and 4NPC as a function of the simulation time. D) Evolution of the absolute binding free-energy $\Delta G_b^\circ$ for 4NPC as a function of the simulation time (red trace) with the reweighted average (green trace) and the mean fluctuation as standard deviation (green bars)

.

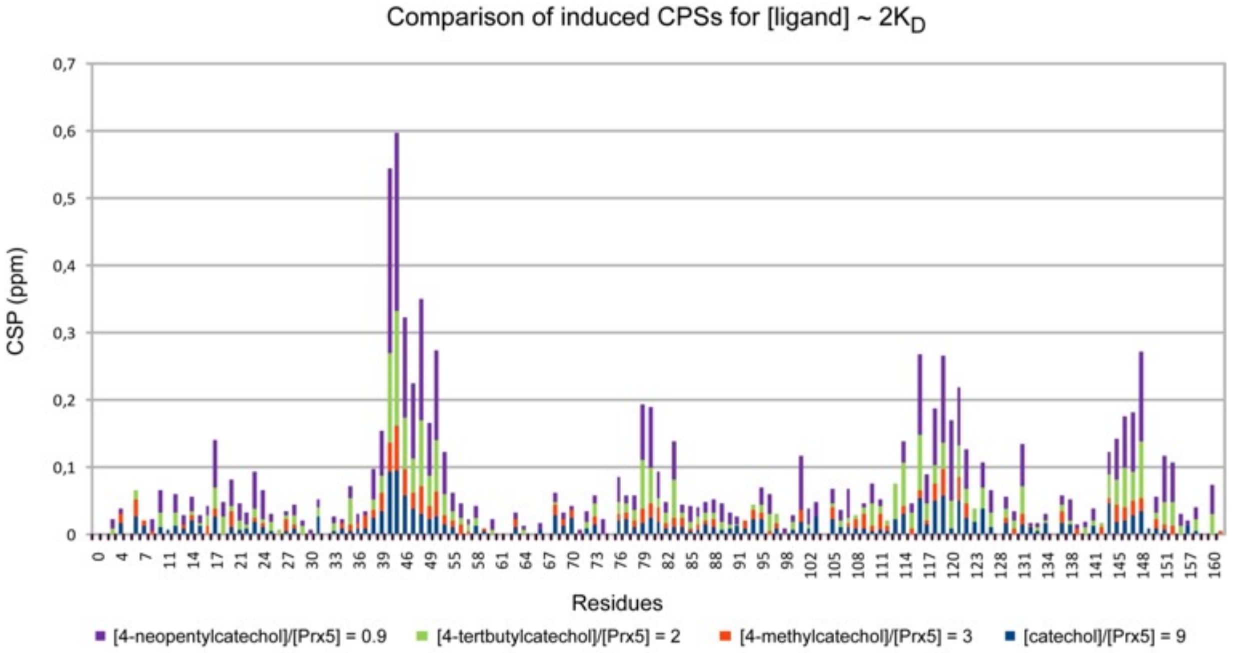


Figure S3: Stacked bar histogram of CSPs induced by catechol derivatives on hPrx5 signals: CAT (bue), 4MEC (orange), 4TBC (green) and 4NPC (purple). Missing data due to spectral overlaps or non-observed amino acids are not represented.

**A**

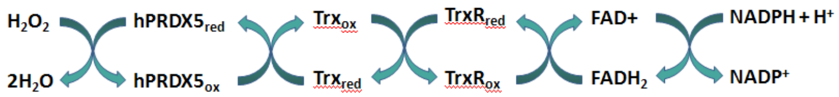


**B**

| Ligand | CAT | 4MEC | 4TBC | 4NPC |
|---|---|---|---|---|
| $IC_{50}$value | 3660 | 801 | 252 | 40 |
| Standard Error | 264 | 163 | 68 | 5 |

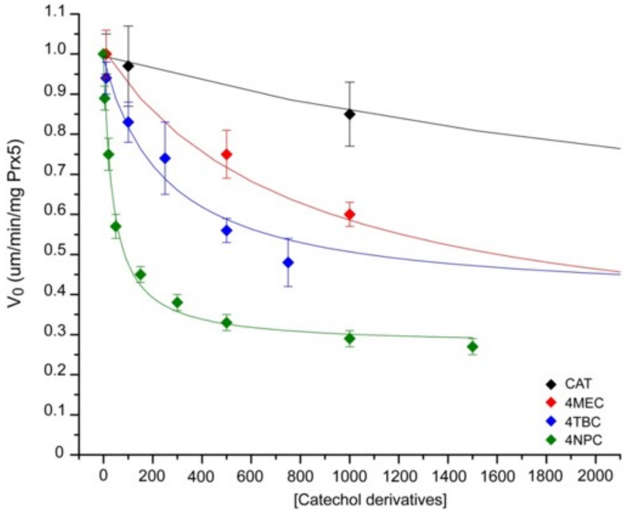


Figure S4: A) Inhibition activity of 4NPC was measured indirectly by the coupled reaction between thioredoxin (Trx) and thioredoxin reductase (TrxR) through the oxidation of NADPH measuring the absorption at 340 nm. (See detailed Methods). B) IC50 results for catechol derivatives and the resulting graph plotting IC50 mean curve measured over 5 separated experiments with error bars represented as standard error. Reaction rates are plotted against catechol derivatives concentration, and fitted with the following non linear regression equation: $y = C + AB/Ax$ with $A$ = IC50 value; $B$ = x intercept at the origin; $C$ = residual activity. The concentration range was evaluated accordingly to the affinity of the ligand for hPrx5. The plot was centered on 4NPC inhibition, so on a range from 0 to 1.5 mM, when 4MEC (0 to 15 mM), 4TBC (0 to 5 mM) and CAT (0 to 80 mM) were analyzed on wider range and are not fully visible on this graph.

**A**

Partial mixed non competitive inhibition model

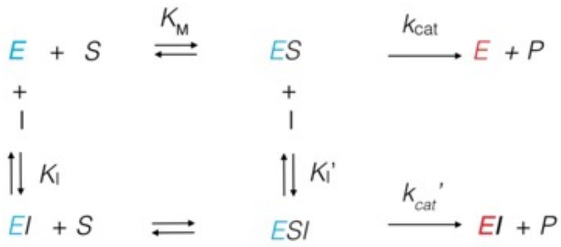


**B**

Partial mixed non competitive

Mixed competitive

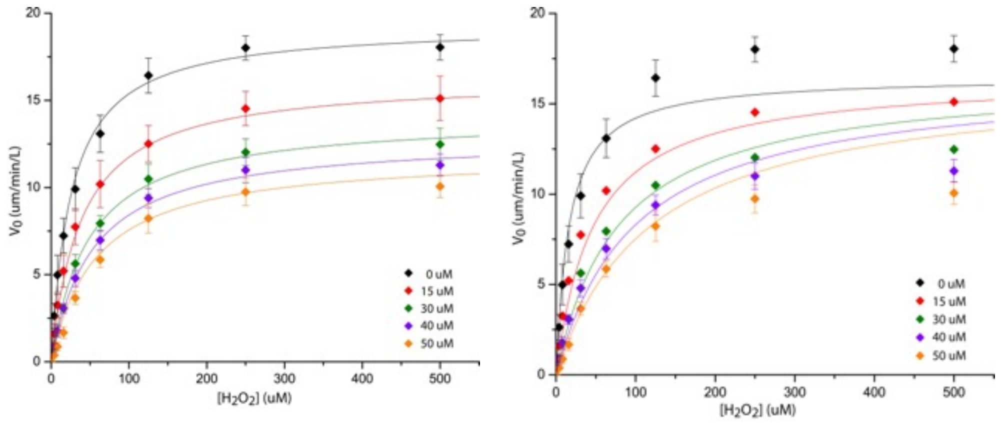


Figure S5: A) Illustration of the partial-mixed type non competitive inhibition model. Two pathways are possible for the inhibition of hPrx5 (E) by 4NPC (I). 4NPC can bind to the free enzyme or to the Michaelis-Menten complex with two different inhibition constants (KI and KI'). Than, as hPrx5 is not completely inhibited, it can also use two different pathways to produce product (P) and be oxidized (E): directly from the ES complex or from the ESI tertiary complex, with two different rate constants kcat and kcat'. B) Non-linear inhibition modeling of 4NPC ligand fitted with DynaFit to a partial mixed non competitive model and a mixed competitive model (for kcat'=0). Initial rate values are plotted against hydrogen peroxide concentration. Experimental points are represented as the mean value of three separated experiments, and error bars as the standard deviation over those experiments.

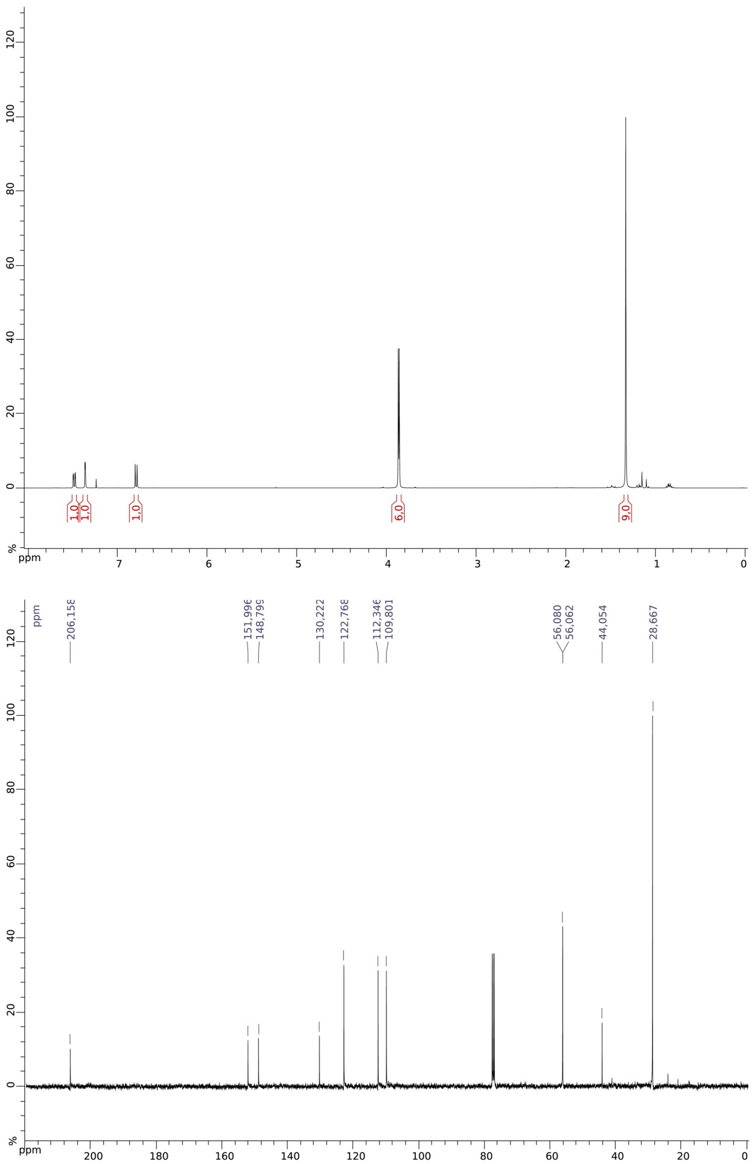


Figure S6: **NMR characterization of the synthesized compounds**. 1-(3,4-dimethoxyphenyl)-2,2-dimethylpropan-1-one (**2**): $^{1}$H NMR(400MHz, $CDCl_3$):1.33(9H, s, $CMe_3$), 3.86 (3H, s, OMe), 3.87 (3H, s, OMe), 6.79 (1H, d, J6-58.5 Hz, H6), 7.36 (1H, d, J3-52 Hz, H3), 7.48 (1H, dd, J5-68.5, J5-32Hz, H5) $^{13}$C NMR (100MHz, $CDCl_3$):28.7 ($CMe_3$), 44.1 ($CMe_3$), 56.06 (OMe), 56.08 (OMe), 109.8 (CHar), 112.3 (CHar), 122.8 (CHar), 130.2 (Cqar), 148.8 (CqarO), 160.0 (CqarO), 206.2 (C=O).

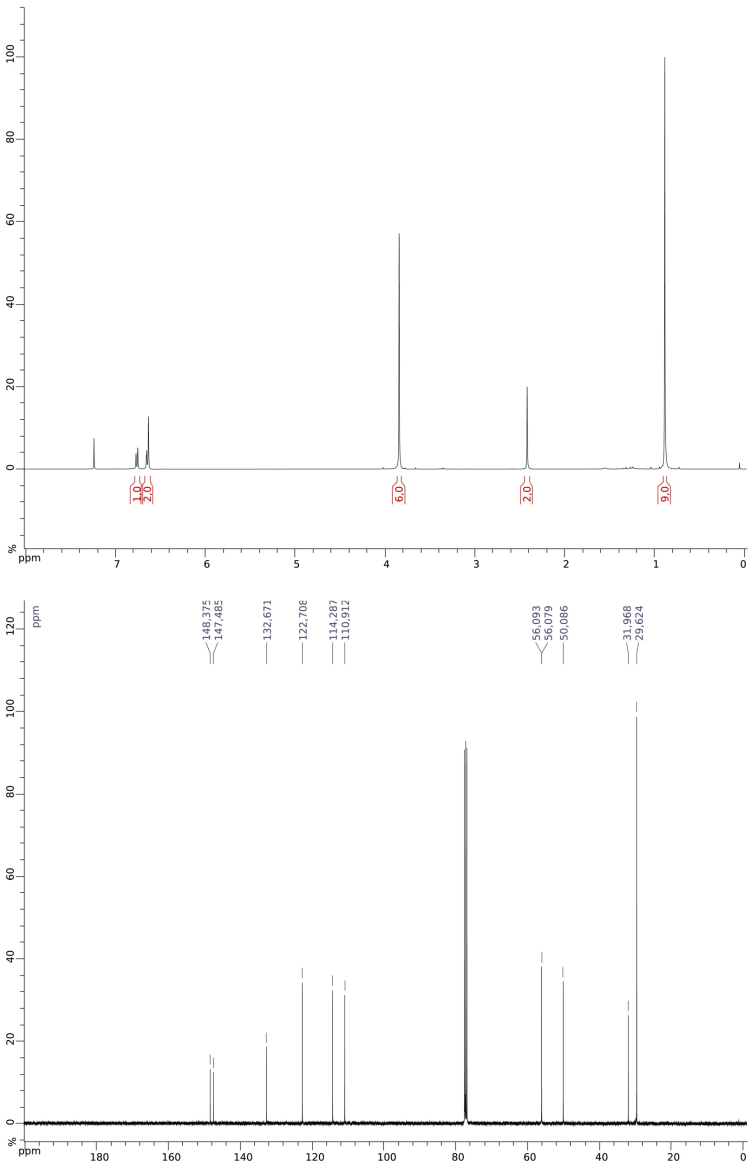


Figure S7: **NMR characterization of the synthesized compounds**. 4-(2,2-Dimethylpropyl)-1,2-dimethoxybenzene (**3**) : $^{1}$H NMR (400MHz, $CDCl_3$): 0.89(9H, s, $CMe_3$), 2.42 (2H, s, $CH_2$), 3.84 (6H, s, OMe), 6.63(1H, s, H3), 6.65(1H, dd, J5-6 8 Hz, J5-3 2 Hz, H5), 6.76 (1H, d,J6-58Hz,H6). $^{13}$C NMR (100MHz, $CDCl_3$): 29.6 ($CMe_3$), 32.0 ($CMe_3$), 50.1 ($CH_2$), 56.08 (OMe), 56.09 (OMe), 110.9 (CHar), 114.3 (CHar), 122.7 (CHar), 132.7 (Cqar), 147.5 (CqarO), 148.4 (CqarO).

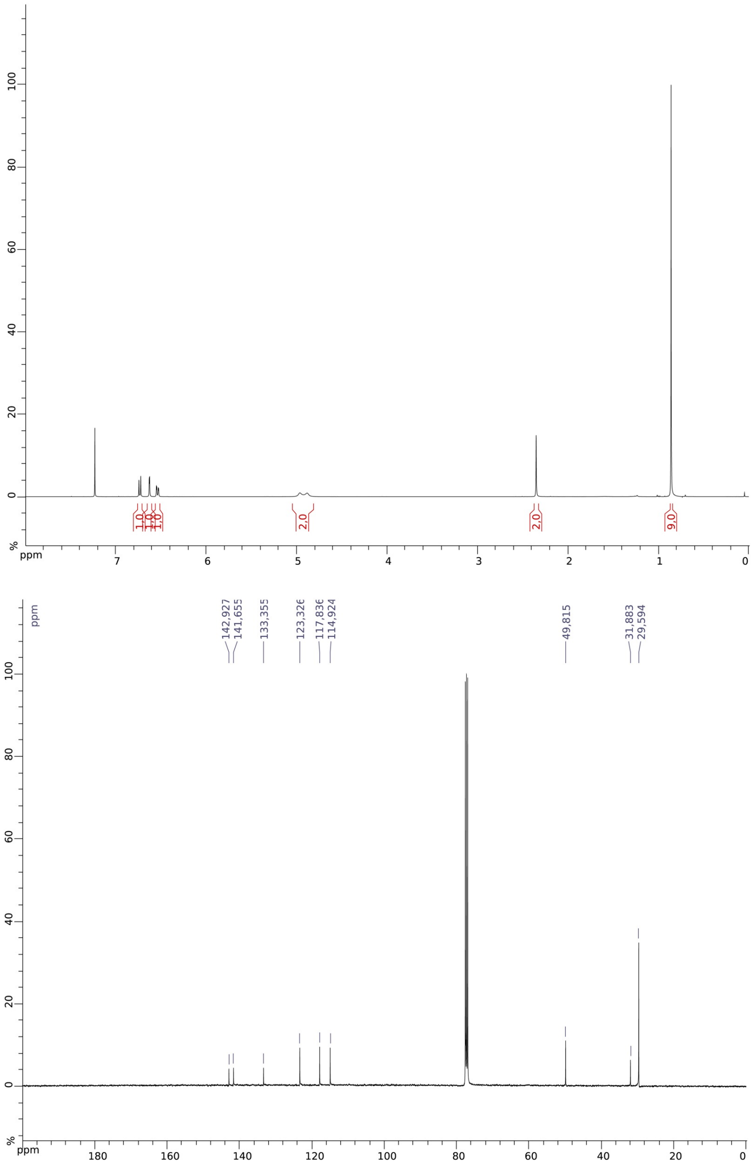


Figure S8: **NMR characterization of the synthesized compounds**. 4-(2,2-Dimethylpropyl)benzene-1,2-diol (**4**) : $^{1}$H NMR (400MHz, $CDCl_3$): 0.86(9H, s, $CMe_3$), 2.35 (2H, s, $CH_2$), 4.88 (1H, s, OH), 4.96 (1H, s, OH),6.54(1H, dd, J5-68 Hz, J5-32 Hz, H5), 6.62 (1H, d, J3-52Hz, H3), 6.73(1H, d, J6-58 Hz, H6). $^{13}$C NMR (100MHz, $CDCl_3$): 29.6 ($CMe_3$), 31.9 ($CMe_3$), 50.0 ($CH_2$), 114.9 (CHar), 117.8 (CHar), 123.3 (CHar), 133.4 (Cqar), 141.7 (CqarO), 142.9 (CqarO).
